# Effects of protein size, thermodynamic stability, and net charge on cotranslational folding on the ribosome

**DOI:** 10.1101/303784

**Authors:** José Arcadio Farías-Rico, Frida Ruud Selin, Ioanna Myronidi, Marie Frühauf, Gunnar von Heijne

## Abstract

During the last five decades, studies of protein folding in dilute buffer solutions have produced a rich picture of this complex process. In the cell, however, proteins can start to fold while still attached to the ribosome (cotranslational folding) and it is not yet clear how the ribosome affects the folding of protein domains of different sizes, thermodynamic stabilities, and net charges. Here, by using arrest peptides as force sensors and on-ribosome pulse proteolysis, we provide a comprehensive picture of how the distance from the peptidyl transferase center in the ribosome at which proteins fold correlates with protein size. Moreover, an analysis of a large collection of mutants of the *E. coli* ribosomal protein S6 shows that the force exerted on the nascent chain by protein folding varies linearly with the thermodynamic stability of the folded state, and that the ribosome environment disfavors folding of domains of high net-negative charge.

## Introduction

More than 40 years ago Anfinsen postulated that “the native structure is determined only by the protein’s amino acid sequence” (1). His work established the protein folding field, which up until recently has focused almost exclusively on biophysical studies of purified proteins that can be reversibly unfolded/refolded *in vitro*. However, it is becoming increasingly clear that many proteins start to fold cotranslationally, *i.e*., while still being synthesized on the ribosome (2-6); elements of secondary structure or small protein domains may even fold completely inside the ribosome exit tunnel (7-13). In contrast to *in vitro* folding, cotranslational folding is still a poorly understood process (14), and we lack basic information such as how protein size and net charge relate to where in the exit tunnel a protein starts to fold, and how protein stability impacts cotranslational folding.

Here, using arrest peptides as force sensors (15, 16) and on-ribosome pulse proteolysis (17), we have analyzed the cotranslational folding of eight protein domains that display cooperative folding *in vitro*. The domains are of different size and fold type, of different thermodynamic stability, and of different net charge. We find direct correlations between protein size and the location in the ribosome exit tunnel at which a protein folds, and between thermodynamic stability and the pulling force generated on the nascent chain during folding. Further, it appears that nascent chain segments with high net-negative charge are pushed out of the negatively charged ribosome exit tunnel before they fold. These findings establish important basic facts about cotranslational folding and reinforce the view of the exit tunnel as an environment that can have a strong impact on protein folding.

## Results

### Folding assays

We have used two assays to follow cotranslational folding of protein domains: an arrest peptide-based assay that makes it possible to detect the tension generated in the nascent chain when a domain folds (8), and an on-ribosome pulse-proteolysis assay where thermolysin resistance is used as an indicator of folding (17).

Translational arrest peptides (APs) are short stretches of polypeptide that interact with the upper parts of the ribosome exit tunnel in such a way that they stall translation when the ribosome reaches the last codon in the AP (18). APs are sensitive to external forces pulling on the nascent chain (19), and stalling efficiency is reduced in proportion to the external pulling force (20, 21). Hence, APs can be used as force sensors to report on cotranslational processes such as protein translocation (22), membrane protein biogenesis (21, 23), and protein folding (8, 15, 20, 24).

The basic construct used in the arrest-peptide cotranslational folding assay is composed of the following elements, Fig. 1a: (*i*) the protein domain to be studied, (*ii*) a variable-length stretch of alternating glycine and serine (GS) residues, (*iii*) the AP, and (*iv*) a 23-residue C-terminal extension that makes it possible to separate arrested and full-length protein products by SDS-PAGE. We define linker length *L* as the number of residues between the protein domain and the end of the AP. As illustrated in Fig. 1b, for small *L* (short linkers) there is not enough space in the ribosome exit tunnel to allow folding at the point during translation when the ribosome reaches the end of the AP, and little force is exerted on the AP. At some intermediate value of *L*, the chain is just long enough to allow folding if stretched, and the force on the AP is increased. At large *L*, the domain is already folded when the ribosome reaches the codon at end of the AP, and again little force is generated. Since APs stall translation less efficiently when under tension (18, 20, 21), the fraction full-length protein (*f_FL_*) produced for constructs with different *L* reflects the variation in force generated on the AP by the folding reaction, and a plot of *f_FL_* vs. *L* can be used to infer where in the exit tunnel folding occurs during translation (15). Mutagenesis studies (16, 25), as well as visualization of folded protein domains located within the exit tunnel by cryo-EM (8, 15, 16, 24) and molecular dynamics simulations (8, 24), show that the dominant peak in a *f_FL_* profile corresponds to folding into the native state (as opposed to, *e.g*., non-specific compaction of the nascent chain), at least for small, single-domain proteins; further support for this notion is provided below. *f_FL_* profiles were recorded for each protein by *in vitro* translation in the PURE^™^ system (26), separation of arrested (*A*) and full-length (*FL*) forms of the protein by SDS-PAGE, and quantitation of the *A* and *FL* bands by phosphoimager analysis.

**Figure 1.**
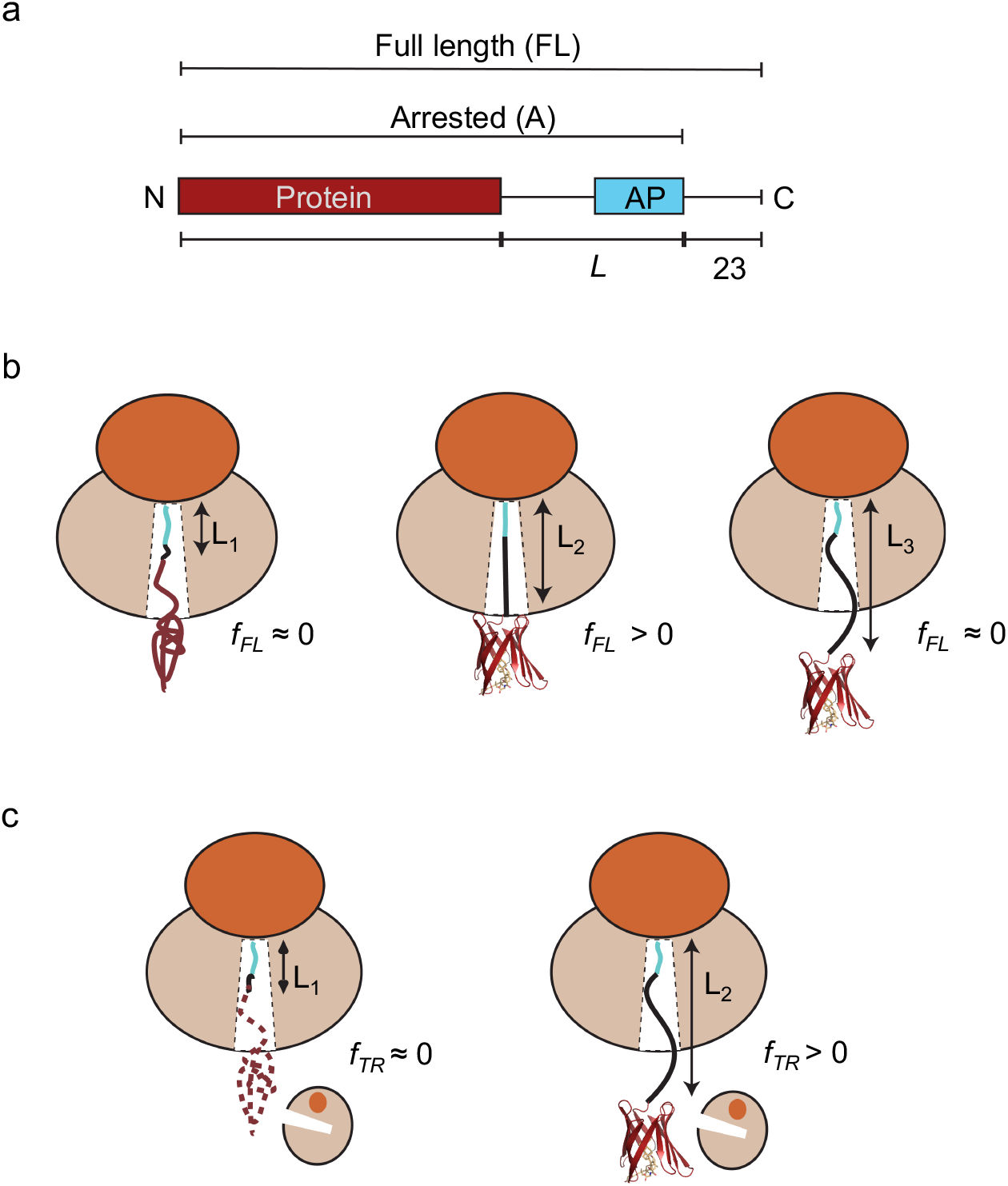
(a) Schematic representation of *in vitro* translated constructs. The protein under study is placed *L* residues upstream of the last proline in the arrest peptide (AP), and a C-terminal 23-residue extension is added downstream of the arrest peptide to ensure that the arrested and full-length forms of the construct can be cleanly separated by SDS-PAGE. (b) The arrest-peptide based assay: at linker length *L_1_*, the protein is too deep in the exit tunnel to be able to fold, hence no force is generated on the AP and mostly arrested nascent chains are produced (*f_FL_* ≈ 0). At *L_2_*, if the linker is stretched beyond its equilibrium length the protein can just reach a location in the exit tunnel where there is sufficient space for it to fold. Some of the folding free energy is therefore stored as elastic energy in the linker, increasing the force on the AP. More full-length proton is produced (*f_FL_* > 0). At *L_3_*, finally, the protein is already folded when the ribosome reaches the C-terminal end of the AP, and little force is exerted on the AP (*f_FL_* ≈ 0). (c) The on-ribosome pulse proteolysis assay: at *L_1_*, the protein is located too deep in the exit tunnel to be able to fold and the nascent chain is hence degraded by a brief thermolysin pulse. At *L_2_*, the protein is folded and hence resistant to proteolysis. Note that the protein is attached to the ribosome via a (GS)_n_ linker that in itself is insensitive to thermolysin.

The pulse-proteolysis assay (17) is based on the premise that a short exposure of a stalled ribosome-nascent chain complex to thermolysin will lead to degradation of the protein if it is partly or wholly unfolded, but not if it is folded, Fig. 1c. Since thermolysin cleaves only on the N-terminal side of hydrophobic residues (27), the (GS)n linker will not be digested. Like the arrest-peptide assay, pulse proteolysis reports on folding into the native state rather than non-specific compaction of the nascent chain (17, 28). Examples of SDS-PAGE gels for both assays are shown in Supplementary Fig. S1.

### The onset of cotranslational folding correlates with domain size

In order to put the possible relation between protein size and the location in the ribosome exit tunnel where folding commences (4, 7) on a firm quantitative footing, we selected eight protein domains of different folds and with sizes ranging from 28 to 128 residues for study (Supplementary Tables 1, 2): the *de novo* designed protein 1FSD (28 residues) (29), a WW domain (35 residues) (30), the *de novo* designed protein EHEE_rd2_0005 (40 residues) (31), protein G (56 residues) (32), a calmodulin fragment (68 residues) (33), ribosomal protein S6 (101 residues) (34), superoxide dismutase 1 (SOD1, 110 residues) (35), and Ileal binding protein (ILBP, 128 residues) (36). We used the moderately strong 17-residue *Escherichia coli* SecM(Ec) AP for all eight proteins (37); in addition we also used the stronger eight-residue SecM(Ms) AP from *Mannheimia succiniciproducens* (38) for protein S6. All *f_FL_* profiles are shown in Fig. 2.

**Figure 2.**
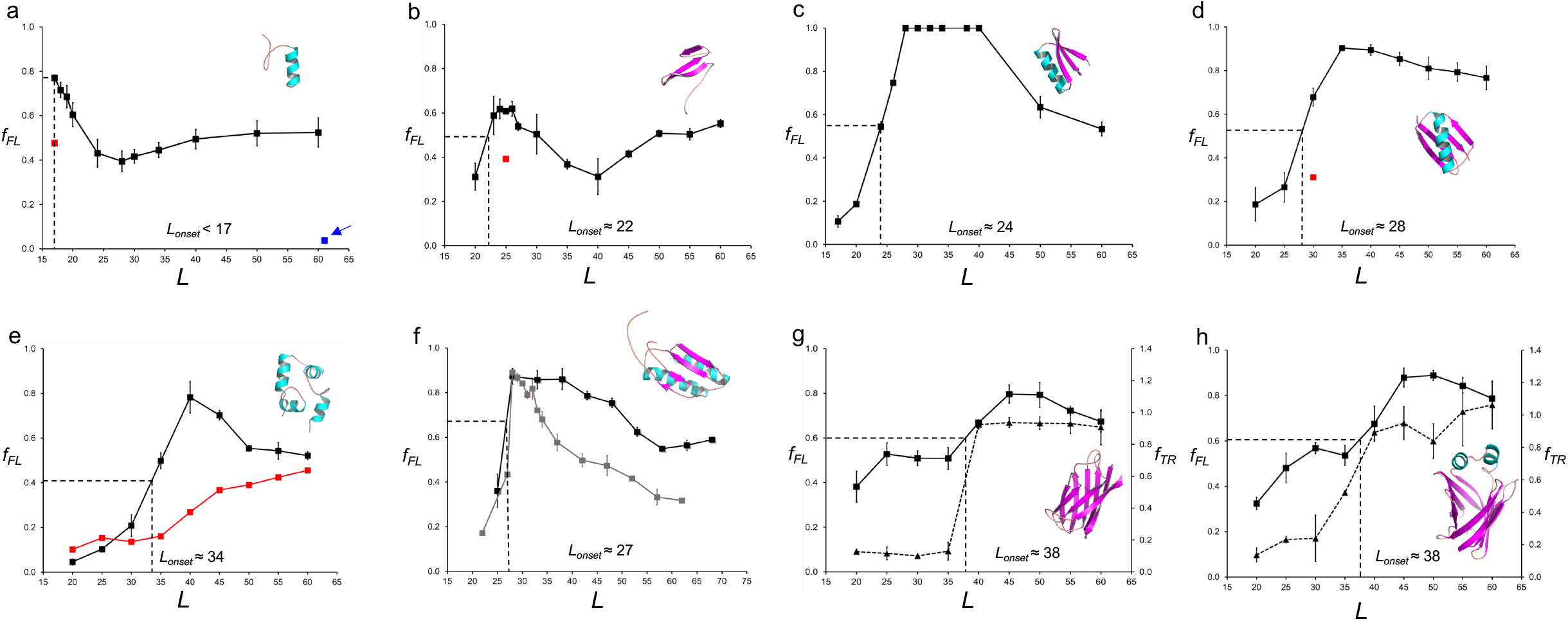
Fraction full-length protein (*f_FL_*) and thermolysin resistant protein (*f_TR_*) plotted as function of *L* for the eight proteins discussed in the text. Each data point is an average of three independent experiments (error bars indicate SEM values). *L_onset_* is the *L* value corresponding to half-maximal peak height (indicated by dotted lines), and *L_max_* is the *L* value corresponding to the maximum of the peak. (a) FSD1. The red data point at *L* = 17 residues is for the destabilizing mutation F21P. The blue data point at *L* = 61 residues (arrow) is for a construct where the (GS)n linker was replaced by a linker derived from the LepB protein (see Methods). (b) WW domain. The red data point at *L* = 25 residues is for the destabilizing mutation Y19A. (c) Designed protein EHEE_rd2_0005. (d) Protein G, the red data point at *L* = 30 residues is for the destabilizing mutation F52L. (e) Calmodulin EF-2-EF-3. The black profile was obtained in 3 mM CaCL_2_, and the red profile in 0.3 mM EGTA. (f) Ribosomal protein S6. The black profile was obtained with the SecM(*Ec*) AP, and the gray profile with the SecM(*Ms*) AP. (g) SOD1 (lacking the metal-loading loops IV and VII). The full curve (squares) is the *f_FL_* profile and the dashed curve (triangles) is the thermolysin-resistance profile (*f_TR_*). (h) ILBP. The full curve (squares) is the *f_FL_* profile and the dashed curve (triangles) is the thermolysin-resistance profile (*f_TR_*).

The *f_FL_* profile for FSD1 (a designed 28-residue αββ zinc finger domain that does not require Zn^2+^ to fold) has a maximum already at *L_max_* = 17 residues, Fig. 2a, significantly different from the *f_FL_* profile previously obtained for the 29-residue zinc finger protein ADR1a that has *L_max_* ≈ 25 residues (8). It has been suggested that the α-helix in FSD1 is more rigid than the β-hairpin, and there is low cooperativity between the a-helix and the β-hairpin during folding (39). Circular dichroism and thermal denaturation suggest that the melting temperature of FSD1 mainly reflects the melting of the a-helix (40). As stretches of a-helix can form deep in the exit tunnel (13, 41, 42), it seems likely that the high *f_FL_* value at *L* = 17 residues reflects the formation of the a-helix rather than folding of the whole domain. To confirm that a folding event is responsible for the peak we introduced a point mutation (F21P) that breaks the a-helix in the *L* = 17 residues construct; this caused *f_FL_* to drop (from 0.77 to 0.47, red data point), as expected.

The WW domain is one of the smallest known β-sheet proteins that can fold autonomously (43). The *f_FL_* profile for the 35-residue WW domain (an antiparallel three-stranded β-sheet) peaks at *L_max_* = 23-25 residues and the protein starts to fold at *L_onset_* ≈ 22 residues, Fig. 2b, in line with previous findings (7). Introduction of the mutation Y19A that reduces the thermodynamic stability of the WW domain (30) caused a reduction of *f_FL_* at *L* = 25 residues from 0.60 to 0.39 (red data point), as expected if the peak reflects a folding event.

The 40-residue protein EHEE_rd2_0005 (βαββ fold) is an artificial mini-protein produced by two rounds of design and selection, and claimed to be “the most stable minimal protein ever found (lacking disulfides or metal coordination)” (31). The peak of the force profile has the highest amplitude of all the tested proteins, with *f_FL_* = 1 between *L* = 28-40 residues. Fig. 2c. Folding starts at *L_onset_* ≈ 24 residues.

The *f_FL_* profile for the 56-residue Protein G (ββαββ fold) (44) has a pronounced peak that starts at *L_onset_* ≈ 28 residues and reaches a maximum at *L_max_* ≈ 35 residues, Fig. 2d. To confirm that the peak is due to folding we introduced the destabilizing mutation F52L (45) that decreased *f_FL_* from 0.7 to 0.3 at *L* = 30 residues (red data point).

Wildtype calmodulin contains four Ca^2+^-binding sites in the so-called EF-hands EF-1 to EF-4. Each EF-hand folds into a helix-loop-helix motif; the 69-residue version of the protein tested here contains only the EF-2 and EF-3 domains (33). This particular version of calmodulin requires Ca^2+^ to fold and has a well-defined hydrophobic core composed mainly of aromatic residues (33). When translated in the presence of 3 mM calcium chloride, calmodulin EF-2-EF-3 displays a peak in the *f_FL_* profile with *L_onset_* ≈ 34 residues and at *L_max_* ≈ 40 residues, Fig. 2e. No comparable peak is seen when the translation reaction is supplemented with the Ca^2+^ chelator EGTA (0.3 mM), validating folding as the source of the peak.

Ribosomal protein S6 from *Thermus thermophylus* is a 101-residue βαββαβ protein with a ferredoxin-like fold (46). It has been subjected to multiple *in vitro* folding studies (47-52). The *f_FL_* profile of S6 has a peak with *L_onset_* ≈ 27 residues and *L_max_* ≈ 28 residues, Fig. 2f. We also obtained an *f_FL_* profile with the stronger SecM(*Ms*) AP and determined the *f_FL_* profile at single-residue resolution around *L_max_*. With the stronger AP, the peak in the *f_FL_* profile is better defined, with *L_onset_* = 28 residues, *L_max_* = 29 residues, and returns to a low baseline of *f_FL_* ≈ 0.3 at long linker lengths.

Super oxide dismutase 1 (SOD1) plays an important role in neurodegenerative disease (53), and its misfolding and aggregation due to de-metalation has been linked to amyotrophic lateral sclerosis. The version of SOD1 tested here has been through a process of loop removal, leaving a β-barrel that does not bind copper or zinc but still folds cooperatively into a globular domain (35). The SOD1 *f_FL_* profile has a major peak at *L_max_* ≈ 45 residues, Fig. 2g. We further studied the folding of SOD1 using pulse proteolysis (17). The protein acquires full thermolysin resistance (as measured by the fraction protease-resistant arrested form of the protein, *f_TR_*) at *L* ≈ 40 residues, coincident with the onset of folding at *L_onset_* ≈ 38 residues seen in the *f_FL_* profile.

The Ileal binding protein (ILBP) is the biggest protein analyzed in this study (128 residues). It shows a significant increase in fraction full length at *L_onset_* ≈ 38 residues and a maximum at *L_max_* = 45-50 residues, Fig. 2h. As was observed with SOD1, thermolysin resistance reaches its maximum coincident with the onset of folding seen in the *f_FL_* profile. We also tested a variant of ILBP that is unable to fold without ligand (36), in which the two helices that cap the β-barrel were removed and replaced by a small GSGS linker (depicted in Supplementary Fig. S1a). In the absence of ligand, the *f_FL_* profile of this variant remains constant throughout the range *L* = 20-60 residues, Supplementary Fig. S1d (red curve). In contrast, upon ligand binding, the mutant protein displays a similar *f_FL_* profile as the wildtype protein, but with somewhat lower amplitude, Supplementary Fig. S1d (gray curve). Notably, in the presence of ligand the mutant protein has a lower melting temperature (T_m_ = 41.2°C) than the apo-wildtype-ILBP (T_m_ = 60.5 °C) (54), supporting the idea that the amplitude of the *f_FL_* peak correlates with the thermodynamic stability of the protein.

One notable feature of all the *f_FL_* profiles obtained with the SecM(*Ec*) AP in Fig. 2 is the rather high *f_FL_* values for the longest linker lengths compared to previously reported *f_FL_* profiles (8, 15, 24). The most obvious difference between the constructs analyzed here and those analyzed before is that we here use a flexible (GS)_n_ linker (to ensure resistance to thermolysin cleavage) rather than a linker with more variable amino acid composition derived from the *E. coli* LepB protein. We therefore substituted the (GS)n linker with the previously used LepB-derived linker in the 1FSD[*L*=61] construct. Indeed, *f_FL_* was reduced to 0.03, much lower than the value of ~0.5 seen for large *L* values with the (GS)_n_ linker, Fig. 2a (blue arrow). The relatively high limiting *f_FL_* values at large *L* values are thus caused by the linker. As seen in the *f_FL_* profile for calmodulin EF-2-EF-3 obtained in the absence of Ca^2+^, Fig. 2e (red curve), the rise in *f_FL_* values caused by the linker becomes noticeable at *L* ≈ 40 residues (when the (GS)_n_ segment is ~25 residues long), and hence does not influence the calculated *L_onset_* values (that are all ≤ 40 residues). An increase in *f_FL_* values at a similar linker length is evident also in Fig. 2a, b.

In summary, we find strong correlations between protein size and the distance from the peptidyl transferase center at which the protein folds, and between the onset of folding as determined from *f_FL_* and *f_TR_* profiles.

### *f_FL_* correlates with thermodynamic stability

If *f_FL_* profiles indeed reflect cotranslational folding to the native state, their amplitude at *L* = *L_max_* should reflect the thermodynamic stability of the folded state. The S6 protein provides a good model system to perform this consistency check, since both thermodynamic stability (Δ*G_D-N_*) and folding/unfolding rate constants (*k_f_*, *k_u_*) have been measured for a large number of point mutants (51). We therefore measured *f_FL_* at linker length *L* = 30 residues, *i.e*., at the middle of the peak of the *f_FL_* profile for wildtype S6 obtained with the strong SecM(*Ms*) AP, for the 16 S6 mutants listed in Supplementary Table 3. As seen in Fig. 3, there is a good correlation between *f_FL_* and *ΔG_D-N_*. A multiple regression analysis including both *ΔG_D-N_* and log *k_f_* values did not yield a statistically significant improvement in the correlation (*p* = 0.063, H0 accepted at the 5% level). Folding kinetics thus does not appear to play a significant role in determining *f_FL_* values under our experimental conditions. However, since the translation rate is ~10-fold slower in PURE^™^ than *in vivo* (55), further studies will be required to fully address this last point.

**Figure 3.**
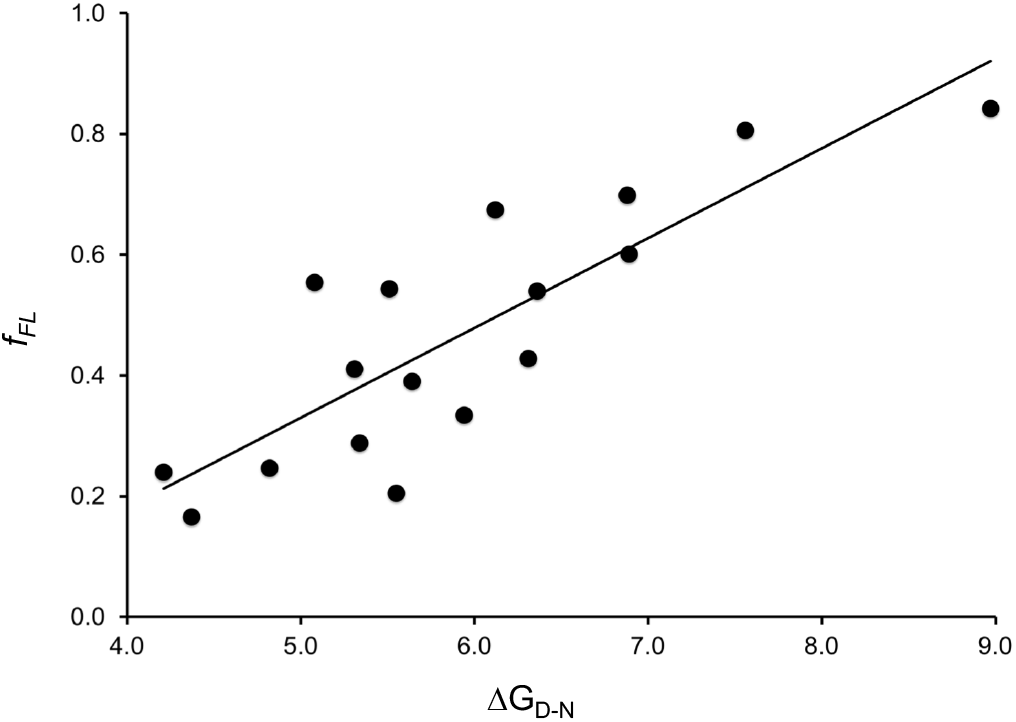
Correlation between *f_FL_* and Δ*G_D-N_* values for wildtype S6 and 16 point mutations listed in Supplementary Table 3. The best-fit linear regression line is shown (*f_FL_* = 0.15 Δ*G_D-N_* - 0.41; *R^2^* = 0.70; *p* = 2·10^−5^). Statistical analysis was performed using the StatPlus:mac Pro software.

The observations that the appearance of the main peak in the S6 (see below), SOD1, and ILBP *f_FL_* profiles correlates with the acquisition of thermolysin resistance in the on-ribosome pulse-proteolysis assay, and that the amplitude of the main peak in the S6 *f_FL_* profile correlates with the thermodynamic stability of the protein, reinforce the notion that the main peak in an *f_FL_* profile represents a *bona fide* folding reaction rather than, *e.g*., the formation of a non-specific collapsed state. The sharp onset of the main peak observed for most proteins analyzed in Fig. 2 and in earlier studies (8, 24, 25) provides further support that *f_FL_* profiles reflect cooperative folding to the native state.

### Net charge and cotranslational folding

Finally, in order to determine how the net charge of a folded protein domain affects cotranslational folding on the ribosome, we compared the *f_FL_* profiles for wildtype S6 and four mutants of different net charge, three of which have previously been characterized *in vitro* in terms of *ΔG_D-N_*, log *k_f_*, and log *k_u_* (52), Fig. 4 and Supplementary Table S4. Wildtype S6 contains 16 positively charged and 16 negatively charged residues (zero net charge). The S6^0^ variant also has zero net charge but lacks all charged residues. The *f_FL_* profile of S6^0^ is broadly similar to that of wildtype S6, but the peak is shifted to somewhat larger *L* values, Fig. 4c. The *f_FL_* profile of the positively charged variant S6^+16-9^ is also similar to the profile for wildtype S6, Fig. 4d, except that the peak is sharper. The *in vitro* folding studies (52) have shown that a mutant that lacks all negatively charged residues (S6^+17^) have slightly accelerated folding and unfolding rates and somewhat lower thermodynamic stability compared to wildtype S6, Supplementary Table 4; presumably, the corresponding Δ*G_D-N_*, log *k_f_*, and log *k_u_* values for S6^+16-9^ are even closer to the wildtype S6 values, explaining the similarity between the two *f_FL_* profiles. S6^0^ has a higher unfolding rate than S6^+17^ and a correspondingly lower thermodynamic stability, consistent with the somewhat higher *L_onset_* and lower amplitude of the *f_FL_* peak.

**Figure 4.**
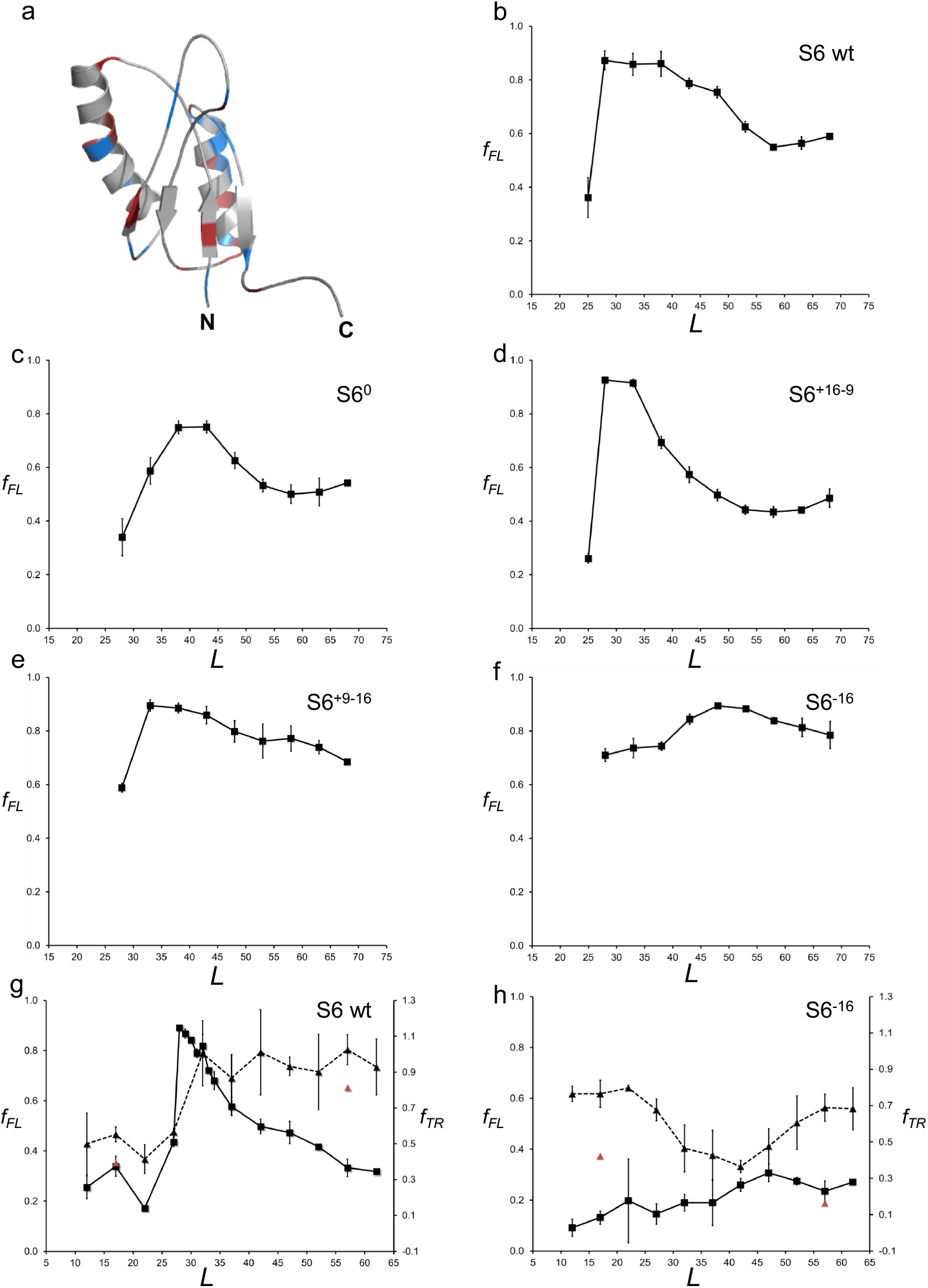
(a) NMR structure of wildtype S6 from *Thermus thermophilus* (PDB 2kjv). Positively charged residues (Arg+Lys) are colored in blue and negatively charged residues (Asp+Glu) in red. (b) *f_FL_* profile of wildtype S6^+16-16^ (for sequences, see Supplementary Figure S2). (c) *f_FL_* profile of S6^0^. (d) *f_FL_* profile of S6^+16-9^. (e) *f_FL_* profile of S6^+9-16^. (f) *f_FL_* profile of S6^-16^. g) *f_FL_* (squares, solid curve) and *f_TR_* (triangles, dashed curve) profiles for wildtype S6. Red data points at *L* = 17 and 57 residues are for the destabilizing mutation L10A. (h) *f_FL_* (squares, solid curve) and *f_TR_* (triangles, dashed curve) profiles for S6^-16^. Red data points at *L* = 17 and 57 residues are for the destabilizing mutation L10A. The SecM(*Ec*) AP was used in panels *a-f* and the SecM(*Ms*) AP in panels *g* and *h*.

In contrast, removal of half of the positive charges from wildtype S6 (variant S6^+9-16^) causes the peak in the *f_FL_* profile to shift from *L_max_* ≈ 28 to *L_max_* ≈ 33 residues, Fig. 4e, and leads to a general increase in *f_FL_* throughout the profile. Removal of the remaining positively charged residues yields the supercharged but not very stable variant S6^-16^; in this case, *f_FL_* ≥ 0.7 for all *L* values, and the profile has a broad maximum around *L_max_* ≈ 50 residues, Fig. 4f.

We also recorded the *f_FL_* and *f_TR_* profiles for wildtype S6 and the supercharged S6^-16^ variant using the stronger SecM(*Ms*) arrest peptide (38). The *f_TR_* profile for wildtype S6 mirrors the *f_FL_* profile, starting to increase at *L* ≈ 27 residues and plateauing at *L_max_* ≈ 32 residues, Fig. 4g. In contrast, the S6^-16^ *f_TR_* profile starts out moderately high at small *L* values, drops to a lower value at *L* = 30-45 residues, and finally increases again to reach a moderately high plateau at *L* > 55 residues, Fig. 5h; the *f_FL_* profile has no distinct peak, similar to the profile obtained with the SecM(Ec) AP but at lower *f_FL_* values (compare Fig. 5f).

**Figure 5.**
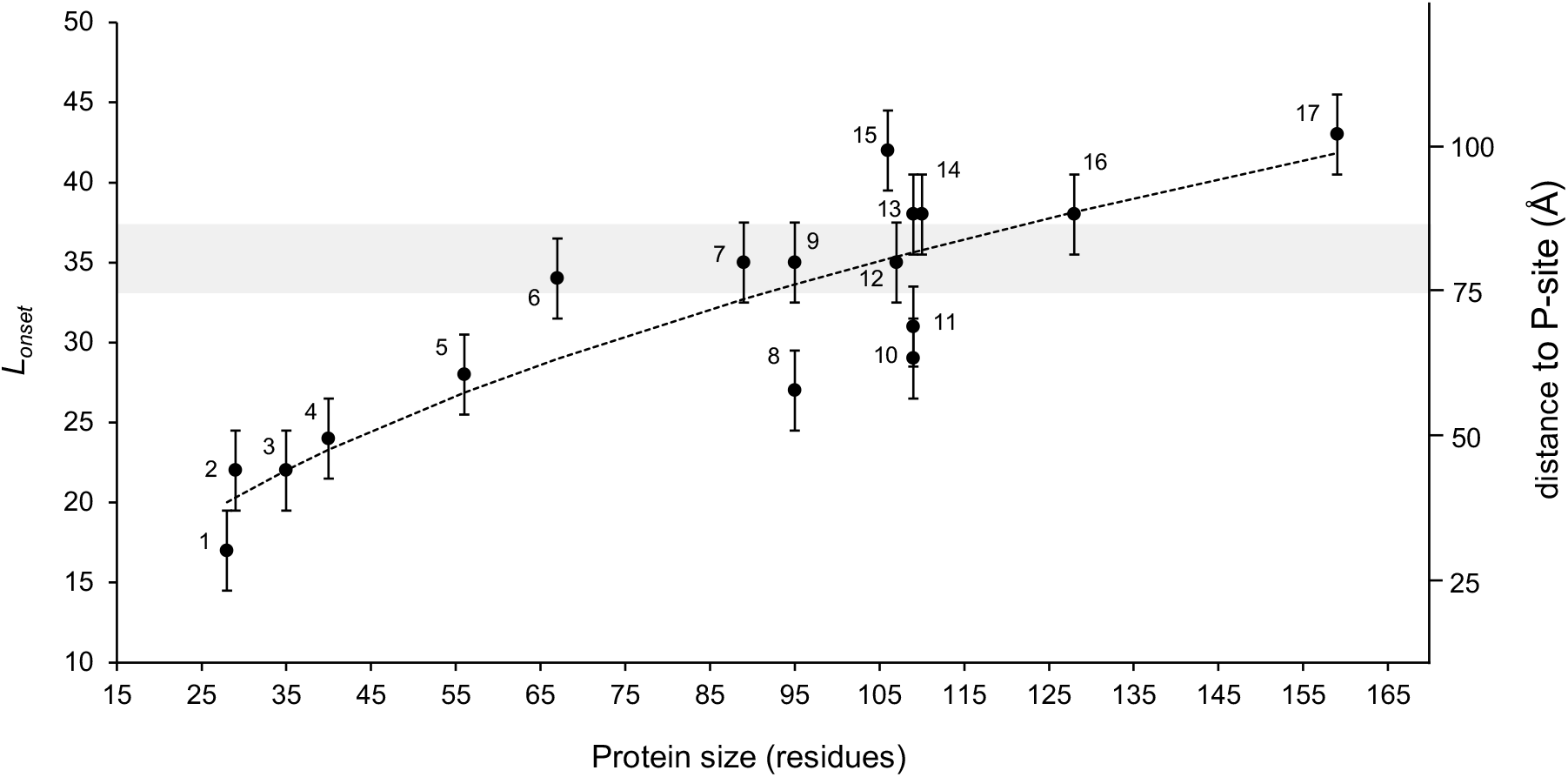
Plot of *L_onset_* vs. proteins size for proteins included in this and previous studies. The shaded gray area indicates the approximate location of the distal end of the exit tunnel, and numbers on the right indicate the approximate distance from the peptidyl transferase center. Error bars of ±2.5 residues have been added to indicate that the *f_FL_* profiles in Fig. 2 were recorded with a 5-residue resolution, hence introducing an uncertainty in the *L_onset_* values. The dotted curve shows the linear best-fit obtained from a log-log plot: *log_10_*(*L_onset_*) = 0.42 (± 0.05) · *log_10_(size*) + 0.69 (± 0.1) (*R*^2^ = 0.80; *p* = 1.4·10^−6^). Proteins are indicated as follows: 1–FSD1, 2–ADR1a1, 3–WW domain, 4–EHEE, 5–Protein G, 6–Calmodulin EF-2-EF-3, 7–Titin I27, 8–S6, 9–TOP7, 10– Spectrin β16, 11– Spectrin R16, 12–PENT, 13–Spectrin R15, 14–SOD, 15–FLN5, 16–ILBP, 17–DHFR (see Supplementary Table 5 for references).

The relatively high thermolysin resistance of S6^-16^ at small *L* values is surprising. To analyze this further, we introduced the destabilizing mutation L10A in both wildtype S6 and S6^-16^ at linker lengths *L* = 17 and *L* = 57 residues. This mutation reduces Δ*G_D-N_* for wildtype S6 from 8.97 to 4.62 kcal/mol (51). The destabilization of S6^-16^ caused by the mutation has not been measured, but the folding free energy of S6^-16^ itself is only 2.89 kcal/mol (52). As seen in Fig. 4g-h (red triangles), *f_TR_* of wildtype S6 is only marginally affected by the mutation—as expected, since even the mutated protein is relatively stable—but is strongly reduced for S6^-16^, at both linker lengths.

We conclude that S6 versions that contain many positively charged residues (wildtype and S6^+17-9^) start to fold well inside the exit tunnel, while S6^0^ and S6^+9-16^ fold near the distal end of the tunnel. For the negatively charged versions S6^+9-16^ and S6^-16^ *f_FL_* values are high for all linker lengths despite their low thermodynamic stability, likely reflecting a strong electrostatic repulsion between the negatively charged exit tunnel and the nascent chain (56-58). There is a hint from the *f_FL_* and *f_TR_* profiles that the supercharged S6^-16^ version may fold outside the exit tunnel, around *L* ≈ 50 residues, but the data are not conclusive on this point.

## Discussion

In this study, we have focused on understanding how three central protein characteristics—size, thermodynamic stability, and net charge—affect cotranslational protein folding.

Our findings on the relation between domain size and the location in the ribosome exit tunnel where folding commences are summarized in Fig. 5, which includes data not only from the present study but also from previous studies on ADR1a (8), three spectrin domains (15), a penta-repeat β-helix protein (25), DHFR (59), the I27 (24) and FNL5 (60) Ig-domain proteins, and the designed protein Top7 (20), (see Supplementary Table 5 for details). From cryo-EM structures of ADR1a (8), spectrin R16 (15), and titin I27 (24) ribosome-nascent chain complexes we know that a ~35 residue linker is required to span the distance between the P-site and a location at the distal end of the exit tunnel, near a prominent loop in the ribosomal protein uL24 (grey shaded area in Fig. 5).

Small domains (≤ 70 residues in size) invariably start to fold while still located inside the tunnel (*L_onset_* < 35 residues), and the smaller they are the deeper in the tunnel they fold. Large domains tend to fold at the distal end of, or even outside, the tunnel (*L_onset_* > 35 residues), but, depending on the location of the first accessible folding nucleus in the structure, may also start to fold while still inside the tunnel, as exemplified by the spectrin domains (15). A linear fit of *log_10_*(*L_onset_*) vs *log*_10_(*size*), Supplementary Fig. S3, shows that *L_onset_* scales approximately with the square root of the protein size over the range covered here (dotted line in Fig. 5). Fig. 5 provides the most complete picture to date of how the gradual increase in tunnel dimensions from the so-called constriction site (~25 Å from the peptidyl transferase site) to the tunnel exit allows protein domains of increasing size to fold.

For the largest proteins analyzed, SOD1 and ILBP, in addition to the main folding peak at *L* ≈ 45-50 residues, we see an early increase in *f_FL_* at *L* ≈ 25-30 residues that might be indicative of a folding intermediate, Fig. 2g-h. A similar increase in *f_FL_* at small *L* values was seen previously for the 189-residue protein DHFR (59). *In vitro*, SOD1 does not seem to fold via an intermediate state, but displays two-state behavior (35). Further studies will be necessary to characterize these putative folding intermediates.

Cotranslational folding of the exceptionally stable 40-residue protein EHEE_rd2_0005 generates sufficient force on the AP to yield *f_FL_* = 1.0 over the whole range *L* = 28-40 residues, Fig. 2c, setting it apart from the other proteins studied here. This and previous observations (16, 21) suggest that the amplitude of the peak in the *f_FL_* profile reflects the thermodynamic stability of the folded protein. Indeed, there is a good correlation between *f_FL_* and Δ*G_D-N_* for a large set of S6 mutants, Fig. 3. Since different proteins fold in different locations inside or outside the exit tunnel, and presumably push against the ribosome in different ways during folding, we do not expect the precise quantitative relation between *f_FL_* and Δ*G_D-N_* found for S6 (see above) to hold for other proteins, but only that the general form of the correlation applies.

Our results for protein S6 variants of different charge content show that protein net charge can have a dramatic effect on the *f_FL_* profile, Fig. 4. In general, the more positive the net charge, the deeper in the exit tunnel S6 folds. We further note that the *f_FL_* profile for the fully uncharged S6^0^ variant is similar to that of wildtype S6, demonstrating that a protein that is totally devoid of charged residues can fold in the exit tunnel. This extends previous results demonstrating efficient folding *in vitro* of a protein completely devoid of charged residues (52, 61) to cotranslational folding.

Strikingly, for the negatively charged variants S6^+9-16^ and S6^-16^, high values of *f_FL_* persist to much larger *L* values than for the neutral and positively charged variants, the peaks in the *f_FL_* profiles are broad and indistinct, and the maximum amplitudes are high, in contrast to what one would expect from the significantly lower thermodynamic stability of S6^-16^ (Supplementary Table 4). Given the net-negative charge of the exit tunnel (56, 58, 62), we hypothesize that the high *f_FL_* values seen for S6^+9-16^ and S6^-16^ are caused by negatively charged residues in the nascent chain being pushed out of the exit tunnel by electrostatic forces even before the protein folds, consistent with ribosome-profiling data showing that stretches of nascent chain of high net-negative charge show reduced ribosome densities (63).

More generally, the fact that domains of up to ~70 residues in size can fold inside the exit tunnel and therefore should be less likely to aggregate or require chaperones for folding might favor the evolution of proteins composed of small (sub)-domains that can initiate early folding inside the ribosome. In support of such a scenario, statistical analyses have identified evolutionarily conserved clusters of rare codons located some 20-50 codons downstream of independently folding sub-domains that may facilitate their cotranslational folding by slowing down translation (64, 65).

Finally, the observation that cotranslational folding generates force on the nascent chain raises the possibility that new sensor systems composed of a ligand binding domain coupled to an arrest peptide may be found in nature, or be constructed for biosensing purposes. In principle, the calmodulin EF-2-EF-3 and ILBP constructs analyzed here could be used as Ca^2+^ and glycochenodeoxycholic acid sensors, respectively, if fused to, *e.g*., GFP (7).

## Acknowledgements

This work was supported by grants from the Knut and Alice Wallenberg Foundation, the Swedish Cancer Foundation, and the Swedish Research Council to GvH. We wish to thank Mikael Oliveberg, Jens Danielsson, Setareh Arya, and Domhnall Iain Henderson for discussions.

## Author contributions

JF and GvH conceived the project. JF and GvH designed the experiments, which were performed by JF, FRS, IM and MF. JF and GvH wrote the manuscript.

## Materials and Methods

### Enzymes and chemicals

Enzymes were purchased from Thermo Scientific (Waltham, MA, USA) and New England Biolabs (Ipswich, MA, USA). Thermolysin from *Geobacillus stearothermophilus* was purchased from Sigma-Aldrich (St. Louis, MO, USA). Oligonucleotides were purchased from Eurofins MWG Operon (Ebersberg, Germany). DNA/RNA purification kits were obtained from Qiagen (Hilden, Germany). The New England Biolabs PURExpress^®^ In Vitro Protein Synthesis Kit was purchased from BioNordika (Stockholm, Sweden). [^35^S]-methionine was purchased from PerkinElmer (Waltham, MA, USA). All other reagents were from Sigma-Aldrich (St. Louis, MO, USA).

### DNA manipulations

The genes of the full sequence design (FSD-1), Pin1 WW domain (*Homo sapiens*), and EHEE design were assembled from oligonucleotides (66). The gene for protein G (*Streptococcus sp*.) was purchased from Genewiz (South Plainfield, NJ). The wildtype calmodulin gene (*Homo sapiens*) was kindly donated by Sara Linse (Lund University) and modified by an adaptation of the original QuickChange site-directed mutagenesis protocol (67) to obtain the fragment tested in this study. Superoxide dismutase 1 (SOD1, *Homo sapiens*) gene (35) was kindly donated by Mikael Oliveberg (Stockholm University). The gene for the Ileal Binding protein (ILBP) from rabbit (*Oryctolagus cuniculus*) was purchased from Invitrogen (Carlsbad, California, United States) and was truncated by an adaptation of the original QuickChange site-directed mutagenesis protocol (67).

The gene for ribosomal protein S6 from *Thermus thermophiles*, along with the genes for the five of its mutants (52), were provided by Mikael Oliveberg.

The constructs were cloned in a pET19b vector under the control of a T7 promoter. As shown in Supplementary Table 2, the small domain (FSD-1, Pin1 WW domain and EHEE) constructs contained a sequence corresponding to an N-terminal unstructured fragment of 158 residues from *E. coli* LepB (21) (to facilitate visualization by SDS/PAGE), a sequence corresponding to a (GS)_n_ flexible linker of 43 or 52 residues (to reach a 60-residues long tether length together with the AP), the sequence corresponding to the SecM AP from *E. coli* (FSTPVWISQAQGIRAGP) (68) (referred here as SecM(*Ec*)) or *M. succiniciproducens* (HAPIRGSP) (69) (referred to here as SecM(*Ms*)) and a sequence corresponding to a 23-residues long LepB-derived C-terminal fragment (to distinguish between full-length and arrested version of proteins after SDS/PAGE). The genes were inserted into the expression vector by Gibson assembly (70). Constructs with increasingly shorter tether length starting from the N-terminal part of the flexible linker and control constructs for the full-length (SecM(*Ec*) FSTPVWISQAQGIRAGA, SecM(*Ms*) HAPIRGSA) and arrested (SecM(*Ec*): FSTPVWISQAQGIRAG*, SecM(*Ms*): HAPIRGS*; where * is a stop codon) forms of the proteins were generated by an adaptation of the original QuickChange site-directed mutagenesis protocol (67). PCR amplified products were subjected to DpnI digestion and purified with GeneJET PCR purification Kit or GeneJET Gel Extraction Kit prior to Gibson assembly. *E. coli* DH5α competent cells were transformed and plated onto LB agar plate supplemented with carbenicillin.

Single colonies were picked to inoculate LB supplemented with ampicillin and were incubated for 12-18 hours at 37°C with shaking. GeneJET Plasmid Miniprep Kit was used for plasmid isolation and purification. All constructs were verified by sequencing (Eurofins Genomics).

### In vitro transcription and translation

Linear constructs were obtained by PCR amplification using oligonucleotides overlapping the T7 promoter and terminator on purified plasmids. After DpnI digestion and purification (GeneJET PCR Purification Kit, Thermo Scientific), the linear constructs were used as template for protein expression. The NEB PURExpress In Vitro Protein Synthesis Kit was used for *in vitro* transcription and translation. The reaction mix was supplemented with [^32^S]-Methionine and the synthesis of the radiolabelled proteins was performed at 37°C, for 15 minutes, under constant shaking at 700 rpm in an Eppendorf 5417R thermomixer. The reaction was stopped by addition of 1:1 volume of 10% ice-cold TCA and the samples were incubated on ice for at least 30 minutes. After centrifugation at 14,000 rpm for 5 minutes at 4°C, the supernatant was discarded and the pellet was re-suspended in sample buffer at 37°C for 15 minutes, under constant shaking at 900 rpm. The samples were treated with RNase I (670 μg/mL) at 37°C for 30 minutes, and were subsequently resolved by SDS/PAGE. All expressions were performed in triplicates.

### On-ribosome pulse-proteolysis

After synthesis of radio-labelled proteins as described in the previous section, the reactions were stopped by the addition of chloramphenicol (instead of TCA) at a final concentration of 3.3 mM. Each sample was divided in two aliquots of equal volume, one for pulse-proteolysis and the other serving as a non-treated control. Pulse-proteolysis was performed by treatment with thermolysin (buffered in 50mM Tris, 500μM ZnCl_2_, pH7.0) at a final concentration of 0.75 mg/m for 1 min. at 37°C, under constant shaking at 700 rpm. The same conditions were applied for the non-treated controls with the difference that thermolysin was not included in the buffer solution. The reaction was stopped by addition of 3μl 500mM EDTA (pH 8.4), and protein precipitation was performed by addition of 1:1 volume of ice-cold 10% TCA and incubation on ice for 30 minutes. As described in the previous section, after centrifugation at 14,000 rpm for 5 minutes, at 4°C, the supernatant was discarded and the pellet was re-suspended in sample buffer at 37°C for 15 minutes, under constant shaking at 900 rpm. The samples were supplemented with RNase I (670 μg/mL) at 37°C, for 30 minutes, and were subsequently resolved by SDS/PAGE. All pulse-proteolysis assays were performed in triplicates.

### Gel electrophoresis and quantitation

Proteins were resolved by SDS-PAGE and visualized on a Fuji FLA-3000 phosphoimager. For *f_FL_* measurements, the bands were quantified to estimate the fraction full-length protein *f_FL_* = *I_FL_*/(*I_FL_* + *I_A_*), where *I_FL_* is the intensity of the band corresponding to the full-length protein and *I_A_* is the intensity of the band corresponding to the arrested form of the protein. *f_FL_*

For pulse proteolysis measurements the thermolysin-resistant fraction resistant was calculated as: *f_TR_* = *I_ATR_* / *I_ABuff_*, where *I_ATR_* is the intensity of the arrested band after pulse proteolysis and *I_ABuff_* is the intensity of the non-treated control band. Bands were quantitated using ImageJ (http://rsb.info.nih.gov/ij/) to obtain an intensity cross section, which was subsequently fit to a Gaussian distribution using in-house software.

### Homology modelling of ILBP mutant

The structure (PBD; 2lba) of chicken Ileal bile acid-binding (ILBP) protein in complex with glycochenodeoxycholic acid (GDCG) was used to produce a homology model (71) of the truncated mutant (ILBP-tm) protein used in this study (59.6 % Seq. Id with 2lba PDB template). We show the estimated position of the ligand (GDCG) in stick representation in Supplementary Fig. 1a.

## Supplementary information

**Supplementary Table 1.**
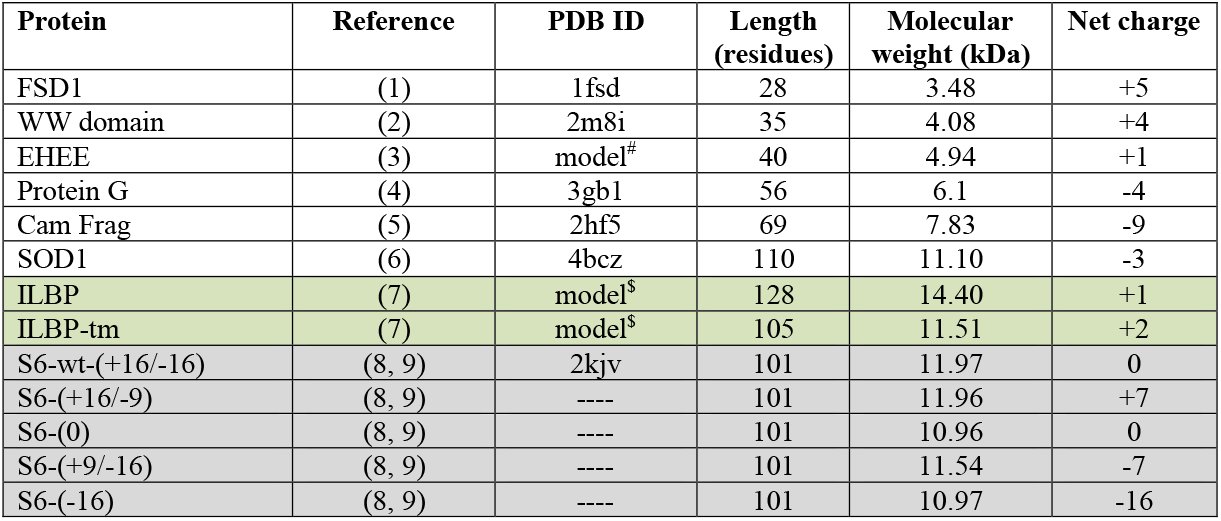
Proteins tested in the study. (#) EHEE is an artificial protein for which only a theoretical 3D model is available. ($) Highlighted in green are the two ILBP proteins used in this study, for which we made homology models based on PDB 2LBA. Highlighted in gray are S6 and four of its variants; there are no structures available for the charge mutants.

**Supplementary Table 2.**
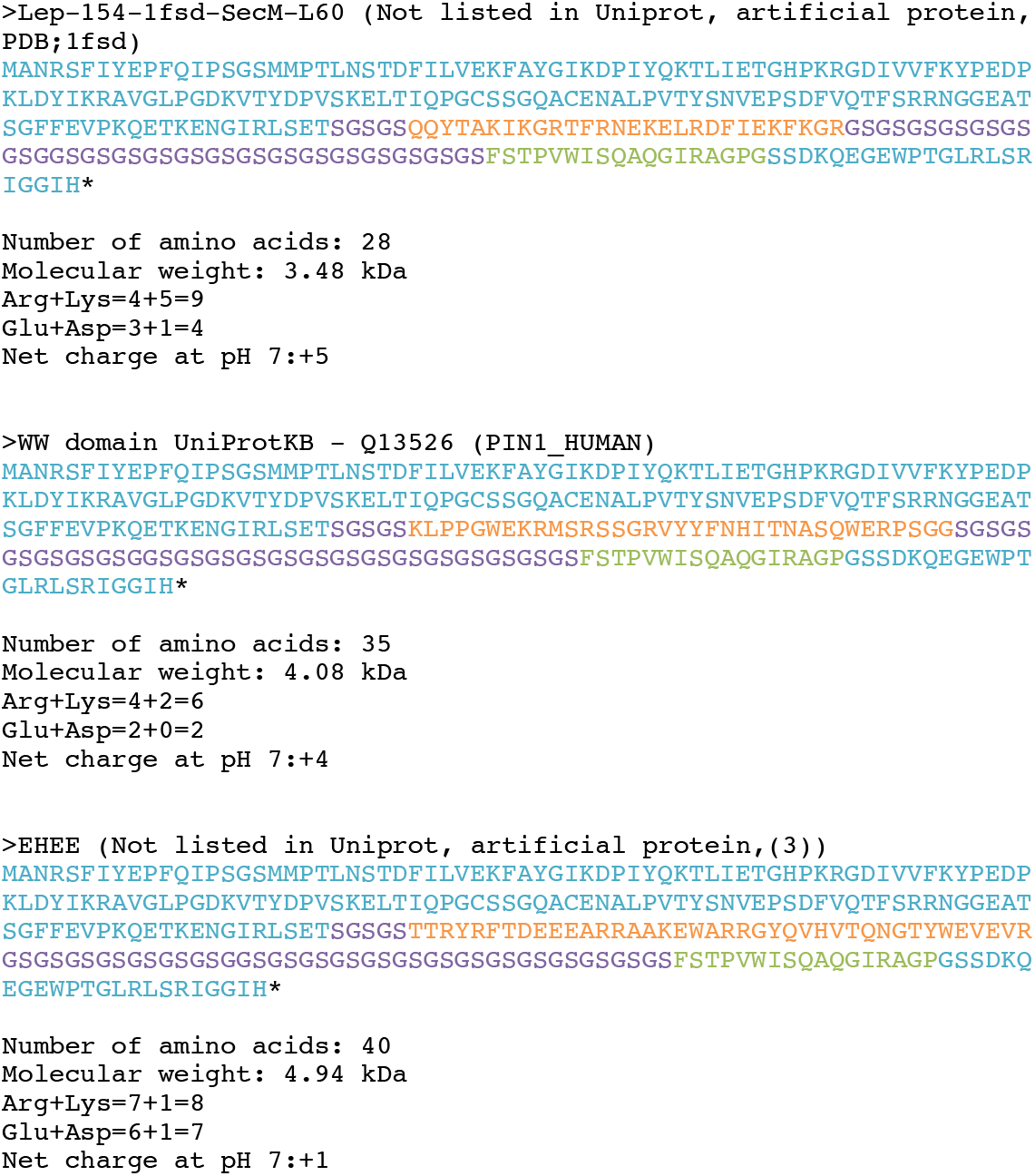
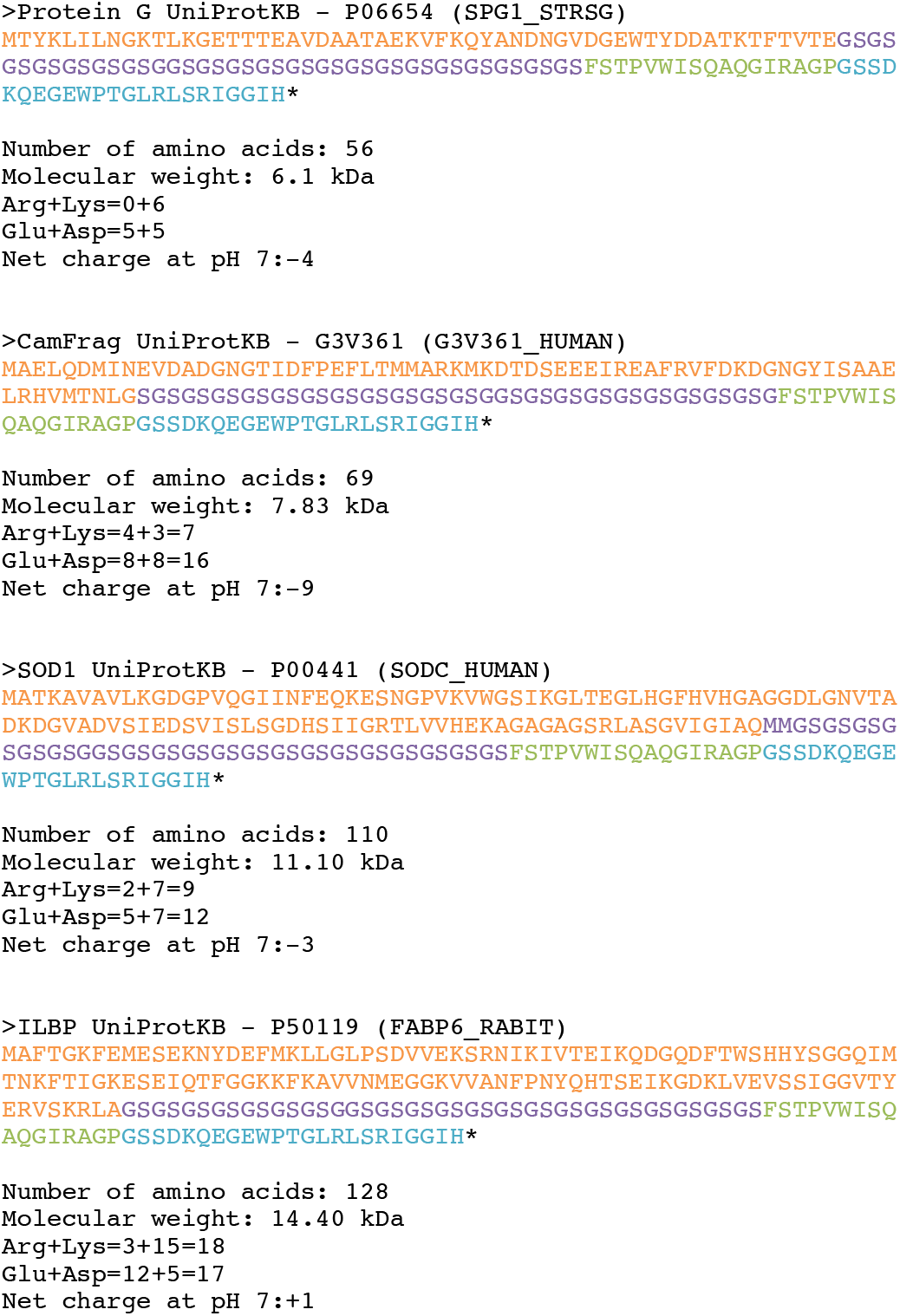
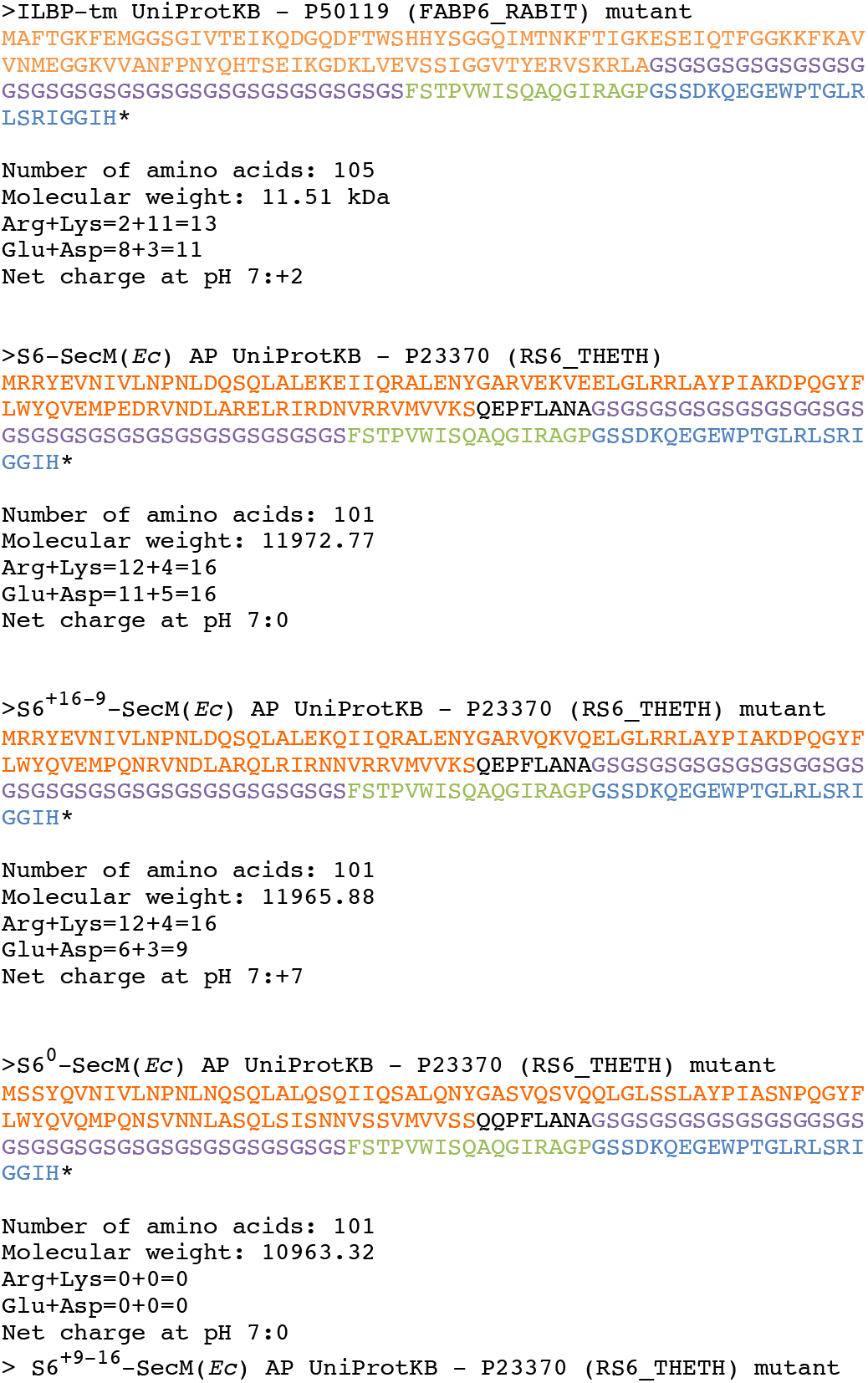
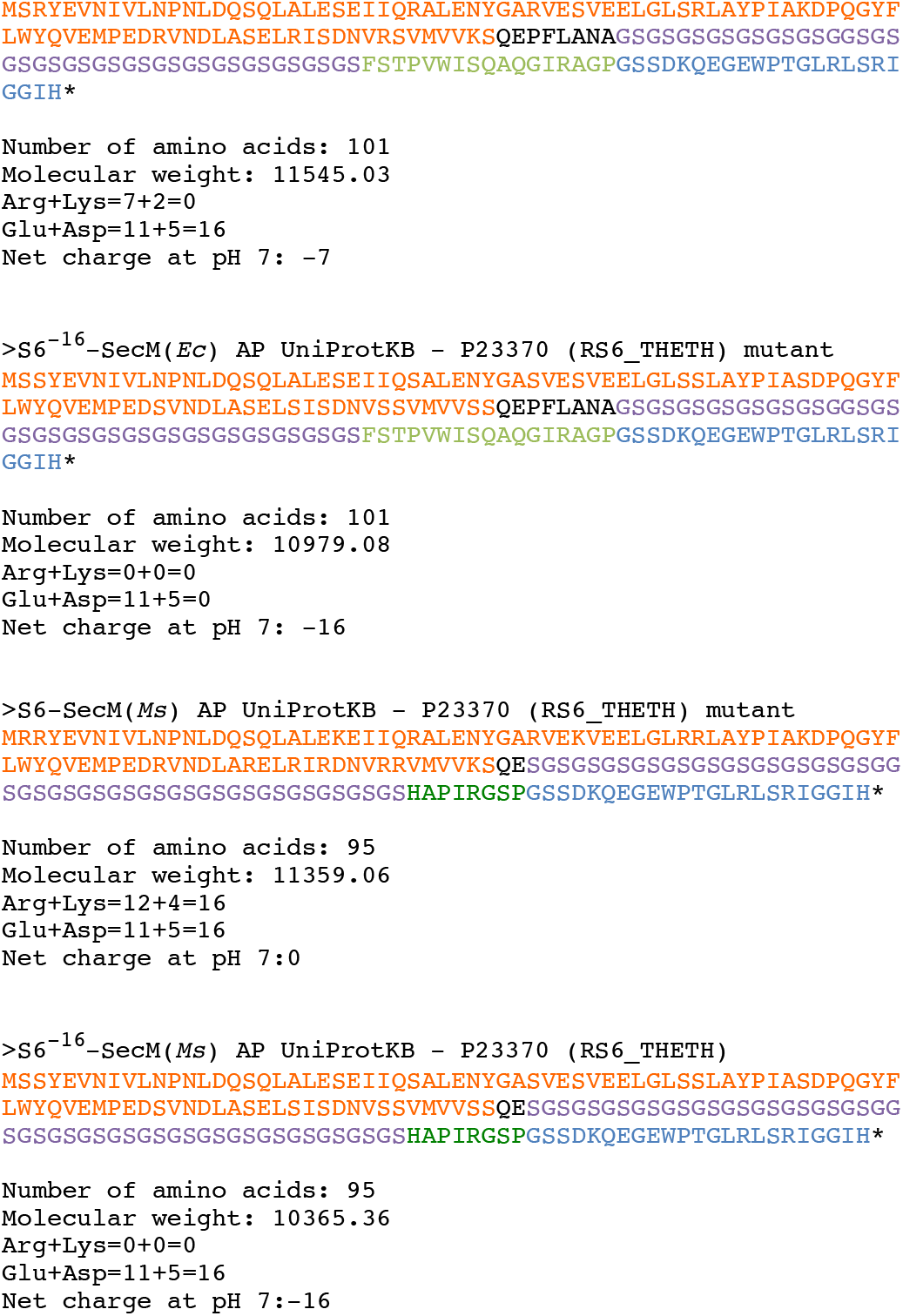
Sequences of the longest construct tested for each protein domain (orange). Shorter constructs were obtained by shortening of the (GS)_n_ linker (purple) For small domains, an unstructured segment from the *E. coli* LepB protein (light blue) was appended to the N terminus in order to make the constructs easy to identify by SDS-PAGE. The arrest peptide (green) is followed by a 23-residue C-terminal tail (light blue), also derived from the LepB protein.

**Supplementary Table 3.** Thermodynamic stabilities (Δ*G_D-N_*) in kcal/mol and *log kf* values (measured at 25 °C in 50 mM MES buffer, pH 6.2) for 16 single-point mutations in protein S6 (10), and fraction full length (*f_FL_*) for the corresponding nascent chain mutants obtained with the SecM(*Ms*) AP at at *L* = 30 residues. The *f_FL_* measurements are averages of 3 independent replicates and their corresponding standard deviations are shown.

| | $\Delta G_{D-N}$ | $\log k_f$ | $f_{FL}$ | ST dev. |
| --- | --- | --- | --- | --- |
| WT | 8.97 | 2.64 | 0.84 | 0.005 |
| V6A | 5.34 | 1.86 | 0.28 | 0.012 |
| I8A | 4.82 | 1.61 | 0.24 | 0.011 |
| L10A | 4.37 | 1.83 | 0.16 | 0.011 |
| L19A | 6.36 | 2.42 | 0.53 | 0.031 |
| I26A | 5.94 | 1.86 | 0.33 | 0.038 |
| L30A | 5.31 | 2.03 | 0.41 | 0.026 |
| V37A | 6.31 | 2.39 | 0.42 | 0.034 |
| L61A | 5.55 | 2.4 | 0.20 | 0.012 |
| Y63A | 5.08 | 2.3 | 0.55 | 0.024 |
| V65A | 5.64 | 2.16 | 0.39 | 0.051 |
| V72A | 7.56 | 2.59 | 0.80 | 0.008 |
| L75A | 6.89 | 2.3 | 0.60 | 0.011 |
| L79A | 4.21 | 2.27 | 0.23 | 0.011 |
| V85A | 5.51 | 2.51 | 0.54 | 0.007 |
| V88A | 6.88 | 2.39 | 0.69 | 0.009 |
| V90A | 6.12 | 2.42 | 0.67 | 0.013 |

**Supplementary Table 4.** Thermodynamic stabilities (Δ*G_D-N_*) in kcal/mol, *log kf*, and *log k_u_* values for S6 variants of different net charge (9). In the *in vitro* studies, the S6^0^ mutant (called S6^+1^ in Ref. 9) was obtained by analyzing the S6^-16^ mutant (called S6^+1-17^ in Ref. 9) at pH 2.3.

| S6 version | $\log k_f$ | $\log k_u$ | $\Delta G_{D-N}$<br>kcal/mol |
| --- | --- | --- | --- |
| WT | 2.53 | -3.51 | 8.24 |
| +17 | 3.34 | -0.86 | 5.72 |
| 0 | 3.28 | 0.35 | 3.99 |
| -16 | 1.48 | -0.64 | 2.89 |

**Supplementary Table 5.**
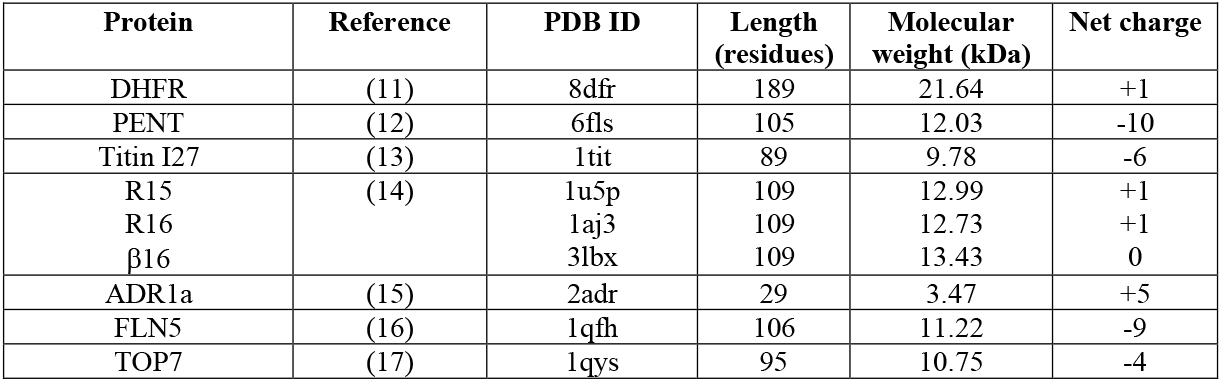
Proteins tested in previous studies

---

**Supplementary Figure S1.**
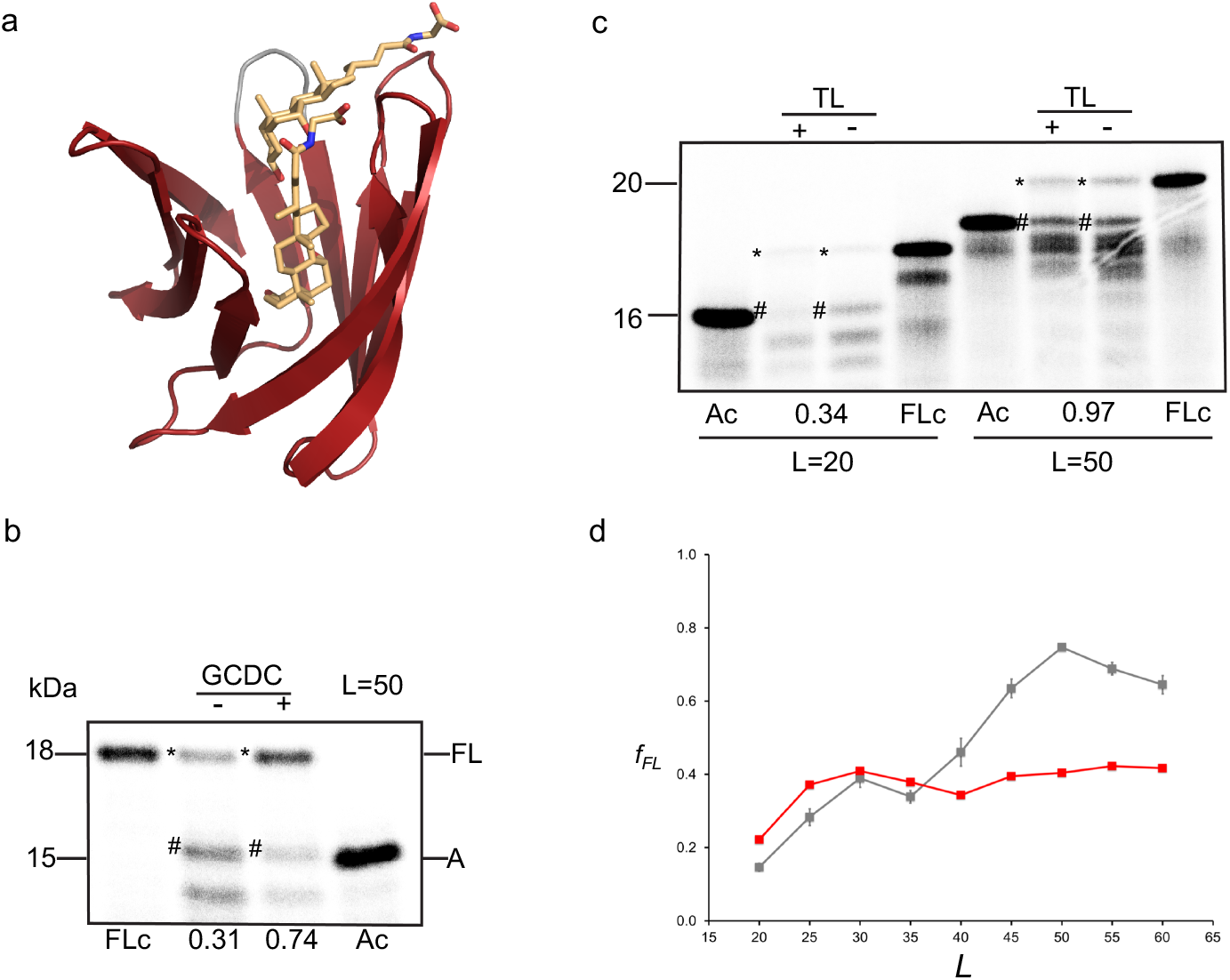
(a) Model for a loop-truncated ILBP mutant used to test the effects of ligand binding on the *f_FL_* profile. The model is based on PDB 2lba, the modeling template showed 60 % sequence identity with the modeled targets and the position of the ligand reflects an estimation of the position in the modeled using as reference the position in the NMR structure. Biochemical data supports the binding of this ligand in all versions of the protein used for this study [17]. (b) *In vitro* translation of the loop-truncated ILBP[L=50] construct with the SecM(Ms) AP, showing that the protein exerts a stronger pulling force on the nascent chain in the presence (+) than in the absence (-) of the ligand GCDC (*f_FL_* = 0.74 *vs*. 0.31; *f_FL_* = *I_FL_*/(*I_FL_*+*I_A_*) where *I_FL_* (or *I_A_*) is the intensity of the band marked * (or #). Full-length (FLc) and arrested (Ac) controls are indicated; the FLc construct has a P-to-A mutation the critical Pro at the end of the AP and does not give any arrested product, the Ac construct has a stop codon inserted directly after the AP. (c) *In vitro* translation and pulse proteolysis of wildtype ILBP at *L* = 20 and *L* = 50 residues. The arrested ILBP nascent chain is susceptible to thermolysin at *L* = 20 residues (*f_TR_* = 0.34) but not at *L* = 50 residues (*f_TR_* = 0.97), compare bands marked # in the ±TL lanes. Full length (FLc) and arrested (Ac) controls are indicated. (d) *f_FL_* profiles for loop-truncated ILBP (red curve) and loop-truncated ILBP translated in the presence of 400 μM of the ligand glycochenodeoxycholic acid (GCDC; grey curve).

**Supplementary Figure S2.**
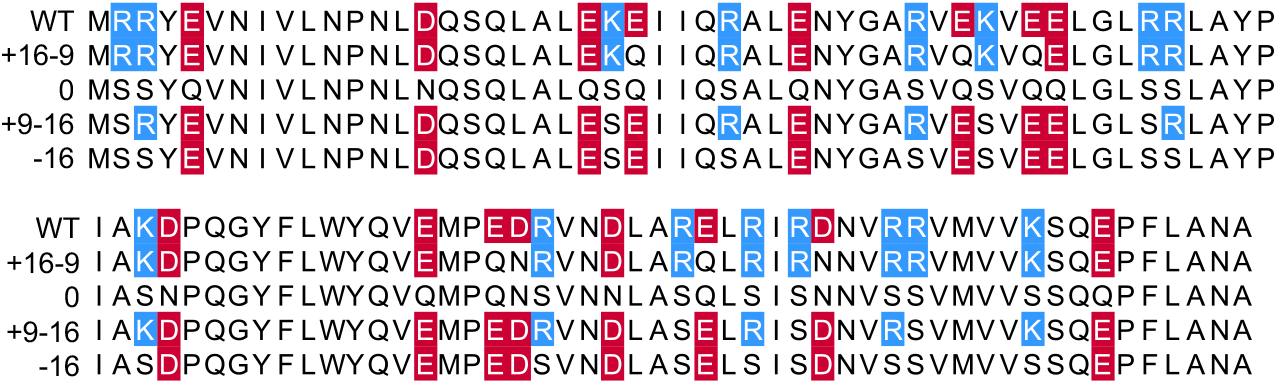
Sequences of the protein S6 charge variants analyzed in Figure 4.

**Supplementary Figure S3.**
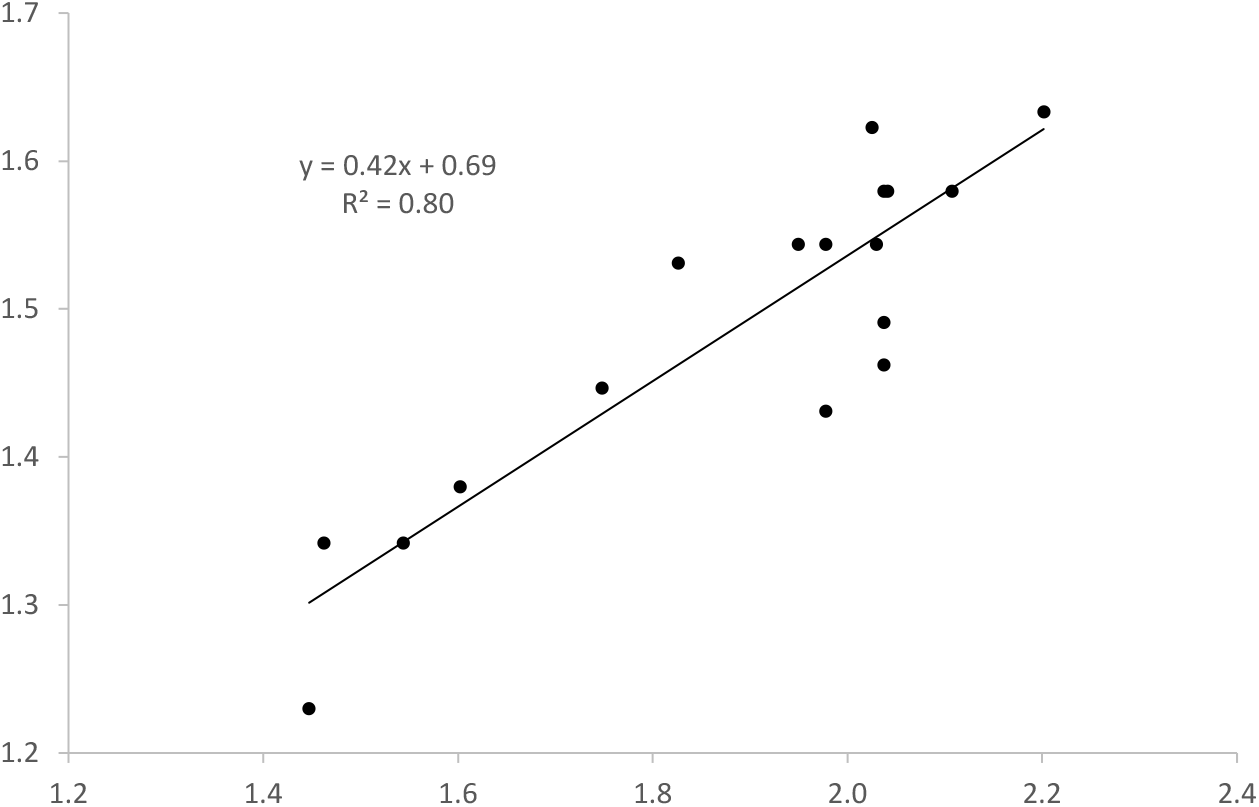
Scatter plot and linear fit of *log_10_*(*L_onset_*) vs *log_10_*(*size*)

## Abbreviations

AP: = arrest peptide

## References

1. Anfinsen CB (1973) Principles that govern the folding of polypeptide chains. Science 181:223–230.

2. Holtkamp W, et al. (2015) Cotranslational protein folding on the ribosome monitored in real time. Science 350:1104–1107.

3. Kim SJ, et al. (2015) Translational tuning optimizes nascent protein folding in cells. Science 348:444–448.

4. Trovato F & O’Brien EP (2016) Insights into Cotranslational Nascent Protein Behavior from Computer Simulations. Annu Rev Biophys 45:345–369.

5. O’Brien EP, Christodoulou J, Vendruscolo M, & Dobson CM (2011) New scenarios of protein folding can occur on the ribosome. J Am Chem Soc 133:513–526.

6. Nicola AV, Chen W, & Helenius A (1999) Co-translational folding of an alphavirus capsid protein in the cytosol of living cells. Nat Cell Biol 1:341–345.

7. Marino J, von Heijne G, & Beckmann R (2016) Small protein domains fold inside the ribosome exit tunnel. FEBS Lett 590:655–660.

8. Nilsson OB, et al. (2015) Cotranslational protein folding inside the ribosome exit tunnel. Cell reports 12:1533–1540.

9. Tu L, Khanna P, & Deutsch C (2014) Transmembrane segments form tertiary hairpins in the folding vestibule of the ribosome. J Mol Biol 426:185–198.

10. Kosolapov A & Deutsch C (2009) Tertiary interactions within the ribosomal exit tunnel. Nat Struct Mol Biol 16:405–411.

11. Conti BJ, Elferich J, Yang Z, Shinde U, & Skach WR (2014) Cotranslational folding inhibits translocation from within the ribosome-Sec61 translocon complex. Nat Struct Mol Biol 21:228–235.

12. Lin PJ, Jongsma CG, Pool MR, & Johnson AE (2011) Polytopic membrane protein folding at L17 in the ribosome tunnel initiates cyclical changes at the translocon. J Cell Biol 195:55–70.

13. Su T, et al. (2017) The force-sensing peptide VemP employs extreme compaction and secondary structure formation to induce ribosomal stalling. eLife 6:e25642.

14. Hingorani KS & Gierasch LM (2014) Comparing protein folding in vitro and in vivo: foldability meets the fitness challenge. Curr Opin Struct Biol 24:81–90.

15. Nilsson OB, et al. (2017) Cotranslational folding of spectrin domains via partially structured states. Nat Struct Mol Biol 24:221–225.

16. Farias-Rico JA, Goetz SK, Marino J, & von Heijne G (2017) Mutational analysis of protein folding inside the ribosome exit tunnel. FEBS Lett 591:155–163.

17. Samelson AJ, Jensen MK, Soto RA, Cate JH, & Marqusee S (2016) Quantitative determination of ribosome nascent chain stability. Proc Natl Acad Sci U S A 113:13402–13407.

18. Ito K & Chiba S (2013) Arrest peptides: cis-acting modulators of translation. Annu Rev Biochem 82:171–202.

19. Butkus ME, Prundeanu LB, & Oliver DB (2003) Translocon “pulling” of nascent SecM controls the duration of its translational pause and secretion-responsive secA regulation. J Bacteriol 185:6719–6722.

20. Goldman DH, et al. (2015) Mechanical force releases nascent chain-mediated ribosome arrest *in vitro* and *in vivo*. Science 348:457–460.

21. Ismail N, Hedman R, Schiller N, & von Heijne G (2012) A biphasic pulling force acts on transmembrane helices during translocon-mediated membrane integration. Nature Struct Molec Biol 19:1018–1022.

22. Ismail N, Hedman R, Lindén M, & von Heijne G (2015) Charge-driven dynamics of nascent-chain movement through the SecYEG translocon. Nat Struct Mol Biol 22:145–149.

23. Cymer F & von Heijne G (2013) Cotranslational folding of membrane proteins probed by arrest-peptide-mediated force measurements. Proc Natl Acad Sci U S A 110:14640–14645.

24. Tian P, et al. (2018) The folding pathway of an Ig domain is conserved on and off the ribosome. BioRxiv doi: https://doi.org/10.1101/253013.

25. Notari L, Martinez-Carranza M, Farias-Rico JA, Stenmark P, & von Heijne G (2018) Cotranslational folding of a pentarepeat β-helix protein. bioRxiv https://doi.org/10.1101/255810.

26. Shimizu Y, et al. (2001) Cell-free translation reconstituted with purified components. Nat Biotechnol 19:751–755.

27. Bogachev MI, Kayumov AR, Markelov OA, & Bunde A (2016) Statistical prediction of protein structural, localization and functional properties by the analysis of its fragment mass distributions after proteolytic cleavage. Sci Rep 6:22286.

28. Park C & Marqusee S (2005) Pulse proteolysis: a simple method for quantitative determination of protein stability and ligand binding. Nat Methods 2:207–212.

29. Dahyiat B & Mayo S (1997) De novo protein design: Fully automated sequence selection. Science 278:82–87.

30. Jager M, Nguyen H, Crane JC, Kelly JW, & Gruebele M (2001) The folding mechanism of a β-sheet: the WW domain. J Mol Biol 311:373–393.

31. Rocklin GJ, et al. (2017) Global analysis of protein folding using massively parallel design, synthesis, and testing. Science 357:168–175.

32. Gronenborn AM, et al. (1991) A novel, highly stable fold of the immunoglobulin binding domain of streptococcal protein G. Science 253:657–661.

33. Lakowski TM, Lee GM, Okon M, Reid RE, & McIntosh LP (2007) Calcium-induced folding of a fragment of calmodulin composed of EF-hands 2 and 3. Protein Sci 16:1119–1132.

34. Ohman A, Oman T, & Oliveberg M (2010) Solution structures and backbone dynamics of the ribosomal protein S6 and its permutant P(54–55). Protein Sci 19:183–189.

35. Danielsson J, Kurnik M, Lang L, & Oliveberg M (2011) Cutting off functional loops from homodimeric enzyme superoxide dismutase 1 (SOD1) leaves monomeric β-barrels. J Biol Chem 286:33070–33083.

36. Rea AM, Thurston V, & Searle MS (2009) Mechanism of ligand-induced folding of a natively unfolded helixless variant of rabbit I-BABP. Biochemistry 48:7556–7564.

37. Zhang J, et al. (2015) Mechanisms of ribosome stalling by SecM at multiple elongation steps. eLife 4.

38. Cymer F, Hedman R, Ismail N, & von Heijne G (2015) Exploration of the arrest peptide sequence space reveals arrest-enhanced variants. J Biol Chem 290:10208–10215.

39. Lei H & Duan Y (2004) The role of plastic beta-hairpin and weak hydrophobic core in the stability and unfolding of a full sequence design protein. J Chem Phys 121:12104–12111.

40. Feng JA, Kao J, & Marshall GR (2009) A second look at mini-protein stability: analysis of FSD-1 using circular dichroism, differential scanning calorimetry, and simulations. Biophys J 97:2803–2810.

41. Tu LW & Deutsch C (2010) A folding zone in the ribosomal exit tunnel for Kv1.3 helix formation. J Mol Biol 396:1346–1360.

42. Bhushan S, et al. (2010) α-Helical nascent polypeptide chains visualized within distinct regions of the ribosomal exit tunnel. Nat Struct Mol Biol 17:313–317.

43. Pucheta-Martinez E, et al. (2016) Changes in the folding landscape of the WW domain provide a molecular mechanism for an inherited genetic syndrome. Sci Rep 6:30293.

44. McCallister EL, Alm E, & Baker D (2000) Critical role of beta-hairpin formation in protein G folding. Nat Struct Biol 7:669–673.

45. Gronenborn AM, Frank MK, & Clore GM (1996) Core mutants of the immunoglobulin binding domain of streptococcal protein G: stability and structural integrity. FEBS Lett 398:312–316.

46. Murzin AG, Brenner SE, Hubbard T, & Chothia C (1995) SCOP: a structural classification of proteins database for the investigation of sequences and structures. J Mol Biol 247:536–540.

47. Lindberg M, Tangrot J, & Oliveberg M (2002) Complete change of the protein folding transition state upon circular permutation. Nat Struct Biol 9:818–822.

48. Otzen DE & Oliveberg M (2002) Conformational plasticity in folding of the split beta-alpha-beta protein S6: evidence for burst-phase disruption of the native state. J Mol Biol 317:613–627.

49. Lindberg MO, Haglund E, Hubner IA, Shakhnovich EI, & Oliveberg M (2006) Identification of the minimal protein-folding nucleus through loop-entropy perturbations. Proc Natl Acad Sci U S A 103:4083–4088.

50. Haglund E, et al. (2012) Trimming down a protein structure to its bare foldons: spatial organization of the cooperative unit. J Biol Chem 287:2731–2738.

51. Haglund E, Lindberg MO, & Oliveberg M (2008) Changes of protein folding pathways by circular permutation. Overlapping nuclei promote global cooperativity. J Biol Chem 283:27904–27915.

52. Kurnik M, Hedberg L, Danielsson J, & Oliveberg M (2012) Folding without charges. Proc Natl Acad Sci U S A 109:5705–5710.

53. Rakhit R & Chakrabartty A (2006) Structure, folding, and misfolding of Cu, Zn superoxide dismutase in amyotrophic lateral sclerosis. Biochim Biophys Acta 1762:1025–1037.

54. Kouvatsos N, et al. (2007) Bile acid interactions with rabbit ileal lipid binding protein and an engineered helixless variant reveal novel ligand binding properties of a versatile beta-clam shell protein scaffold. J Mol Biol 371:1365–1377.

55. Capece MC, Kornberg GL, Petrov A, & Puglisi JD (2015) A simple real-time assay for in vitro translation. RNA 21:296–305.

56. Petrone PM, Snow CD, Lucent D, & Pande VS (2008) Side-chain recognition and gating in the ribosome exit tunnel. Proc Natl Acad Sci U S A 105:16549–16554.

57. Knight AM, et al. (2013) Electrostatic effect of the ribosomal surface on nascent polypeptide dynamics. ACS Chem Biol 8:1195–1204.

58. Lu J, Kobertz WR, & Deutsch C (2007) Mapping the electrostatic potential within the ribosomal exit tunnel. J Mol Biol 371:1378–1391.

59. Nilsson OB, Müller-Lucks A, Kramer G, Bukau B, & von Heijne G (2016) Trigger factor reduces the force exerted on the nascent chain by a cotranslationally folding protein. J Mol Biol 428:1356–1364.

60. Cabrita LD, et al. (2016) A structural ensemble of a ribosome-nascent chain complex during cotranslational protein folding. Nat Struct Mol Biol 23:278–285.

61. Hojgaard C, et al. (2016) A Soluble, Folded Protein without Charged Amino Acid Residues. Biochemistry 55:3949–3956.

62. Fedyukina DV, Jennaro TS, & Cavagnero S (2014) Charge segregation and low hydrophobicity are key features of ribosomal proteins from different organisms. J Biol Chem 289:6740–6750.

63. Requiao RD, et al. (2017) Protein charge distribution in proteomes and its impact on translation. PLoS Comput Biol 13:e1005549.

64. Jacobs WM & Shakhnovich EI (2017) Evidence of evolutionary selection for cotranslational folding. Proc Natl Acad Sci U S A 114:11434–11439.

65. Chaney JL, et al. (2017) Widespread position-specific conservation of synonymous rare codons within coding sequences. PLoS Comput Biol 13:e1005531.

66. Rouillard JM, et al. (2004) Gene2Oligo: oligonucleotide design for in vitro gene synthesis. Nucleic Acids Res 32:W176–180.

67. Zheng L, Baumann U, & Reymond JL (2004) An efficient one-step site-directed and site-saturation mutagenesis protocol. Nucleic Acids Res 32:e115.

68. Nakatogawa H & Ito K (2002) The ribosomal exit tunnel functions as a discriminating gate. Cell 108:629–636.

69. Yap MN & Bernstein HD (2009) The plasticity of a translation arrest motif yields insights into nascent polypeptide recognition inside the ribosome tunnel. Mol Cell 34:201–211.

70. Gibson DG, et al. (2009) Enzymatic assembly of DNA molecules up to several hundred kilobases. Nat Methods 6:343–345.

71. Bertoni M, Kiefer F, Biasini M, Bordoli L, & Schwede T (2017) Modeling protein quaternary structure of homo- and hetero-oligomers beyond binary interactions by homology. Sci Rep 7:10480.

## Supplementary References

1. Dahiyat B, Sarisky C, & Mayo S (1997) De novo protein design: towards fully automated sequence selection. J Mol Biol 273:789–796.

2. Luh LM, et al. (2013) Molecular crowding drives active Pin1 into nonspecific complexes with endogenous proteins prior to substrate recognition. J Am Chem Soc 135:13796–13803.

3. Rocklin GJ, et al. (2017) Global analysis of protein folding using massively parallel design, synthesis, and testing. Science 357:168–175.

4. Kuszewski J, Gronenborn AM, & Clore GM (1996) Improving the quality of NMR and crystallographic protein structures by means of a conformational database potential derived from structure databases. Protein Sci 5:1067–1080.

5. Lakowski TM, Lee GM, Okon M, Reid RE, & McIntosh LP (2007) Calcium-induced folding of a fragment of calmodulin composed of EF-hands 2 and 3. Protein Sci 16:1119–1132.

6. Danielsson J, et al. (2013) Global structural motions from the strain of a single hydrogen bond. Proc Natl Acad Sci U S A 110:3829–3834.

7. Rea AM, Thurston V, & Searle MS (2009) Mechanism of ligand-induced folding of a natively unfolded helixless variant of rabbit I-BABP. Biochemistry 48:7556–7564.

8. Ohman A, Oman T, & Oliveberg M (2010) Solution structures and backbone dynamics of the ribosomal protein S6 and its permutant P(54-55). Protein Sci 19:183–189.

9. Kurnik M, Hedberg L, Danielsson J, & Oliveberg M (2012) Folding without charges. Proc Natl Acad Sci U S A 109:5705–5710.

10. Haglund E, Lindberg MO, & Oliveberg M (2008) Changes of protein folding pathways by circular permutation. Overlapping nuclei promote global cooperativity. J Biol Chem 283:27904–27915.

11. Nilsson OB, Müller-Lucks A, Kramer G, Bukau B, & von Heijne G (2016) Trigger factor reduces the force exerted on the nascent chain by a cotranslationally folding protein. J Mol Biol 428:1356–1364.

12. Notari L, Martinez-Carranza M, Farias-Rico JA, Stenmark P, & von Heijne G (2018) Cotranslational folding of a pentarepeat β-helix protein. bioRxiv https://doi.org/10.1101/255810.

13. Tian P, et al. (2018) The folding pathway of an Ig domain is conserved on and off the ribosome. BioRxiv doi: https://doi.org/10.1101/253013.

14. Nilsson OB, et al. (2017) Cotranslational folding of spectrin domains via partially structured states. Nat Struct Mol Biol 24:221–225.

15. Nilsson OB, et al. (2015) Cotranslational protein folding inside the ribosome exit tunnel. Cell reports 12:1533–1540.

16. Cabrita LD, et al. (2016) A structural ensemble of a ribosome-nascent chain complex during cotranslational protein folding. Nat Struct Mol Biol 23:278–285.

17. Goldman DH, et al. (2015) Mechanical force releases nascent chain-mediated ribosome arrest in vitro and in vivo. Science 348:457–460.

